# Gut Microbiome Dynamics and Predictive Value in Hospitalized COVID-19 Patients: A Comparative Analysis of Shallow and Deep Shotgun Sequencing

**DOI:** 10.1101/2023.11.29.568526

**Authors:** Katarzyna Kopera, Tomasz Gromowski, Witold Wydmański, Karolina Skonieczna-Żydecka, Agata Muszyńska, Kinga Zielińska, Anna Wierzbicka-Woś, Mariusz Kaczmarczyk, Roland Kadaj-Lipka, Danuta Cembrowska-Lech, Kornelia Januszkiewicz, Katarzyna Kotfis, Wojciech Witkiewicz, Magdalena Nalewajska, Wiktoria Feret, Wojciech Marlicz, Igor Łoniewski, Paweł P. Łabaj, Grażyna Rydzewska, Tomasz Kosciolek

## Abstract

The COVID-19 pandemic caused by SARS-CoV-2 has led to a wide range of clinical presentations, with respiratory symptoms being common. However, emerging evidence suggests that the gastrointestinal (GI) tract is also affected, with angiotensin-converting enzyme 2, a key receptor for SARS-CoV-2, abundantly expressed in the ileum and colon. The virus has been detected in GI tissues and fecal samples, even in cases with negative respiratory results. GI symptoms have been associated with an increased risk of ICU admission and mortality. The gut microbiome, a complex ecosystem of around 40 trillion bacteria, plays a crucial role in immunological and metabolic pathways. Dysbiosis of the gut microbiota, characterized by a loss of beneficial microbes and decreased microbial diversity, has been observed in COVID-19 patients, potentially contributing to disease severity. We conducted a comprehensive gut microbiome study in 204 hospitalized COVID-19 patients using both shallow and deep shotgun sequencing methods. We aimed to track microbiota composition changes induced by hospitalization, link these alterations to clinical procedures (antibiotics administration) and outcomes (ICU referral, survival), and assess the predictive potential of the gut microbiome for COVID-19 prognosis. Shallow shotgun sequencing was evaluated as a cost-effective diagnostic alternative for clinical settings.

## 1. Introduction

The World Health Organization declared the Coronavirus Disease 2019 (COVID-19), caused by the SARS-CoV-2 coronavirus, to be a pandemic on March 11, 2020. COVID-19 is a respiratory disease with a wide range of clinical appearances. It may manifest as asymptomatic or mild infection with cough and fever to severe pneumonia with multiple organ failure and acute respiratory distress syndrome (Hu et al., 2020).

Besides common pulmonary symptoms of COVID-19, there is data on the infection of the gastrointestinal tract. Angiotensin-converting enzyme 2, a critical receptor during viral entry of SARS-CoV-2 to the host cells, is abundantly expressed in the ileum and colon, especially in differentiated enterocytes (Burgueño et al., 2020). Moreover, SARS-CoV-2 has been found within the tissues of the entire gastrointestinal (GI) tract, and even in cases when reverse transcription polymerase chain reaction results from respiratory samples were negative, a large percentage of patients still shed the virus in their faeces (Chen et al., 2020). Therefore, SARS-CoV-2 infection directly influences the GI tract, presumably acting as an extrapulmonary location for virus activity and reproduction (Wölfel et al., 2020; Zhou et al., 2020a). Interestingly, the GI symptoms were associated with a significantly increased risk of intensive care unit (ICU) admission and mortality (Woodruff et al., 2020).

In the gastrointestinal tract, it is estimated that there are about 40 trillion bacteria that, along with their genes, constitute the gut microbiome (Sender et al., 2016; Thursby and Juge, 2017). Through intricate pathways, the microbiome contributes significantly to the immunological and metabolic pathways, affecting both the etiology of illnesses and health maintenance (Durack and Lynch, 2019). This effect of the microbiome on the course of the disease and health management was demonstrated in COVID-19 patients. Dysbiosis of the gut microbiota, defined as the loss of beneficial microbes, the proliferation of potentially harmful microbes, and decreased microbial diversity, raises levels of the SARS-CoV-2 target angiotensin-converting enzyme 2, which causes epithelial damage and inflammation (Thevaranjan et al., 2018). Moreover, SARS-CoV-2 activates the NLRP3 inflammasome, which triggers a cascade of pro-inflammatory mechanisms (Ratajczak and Kucia, 2020). The gut microbiota can activate or inhibit the NLRP3 inflammasome and thus can condition the strength of inflammasome stimulation during SARS-CoV-2 virus infection (Dang and Marsland, 2019). Intestinal biocenosis has been found to be altered in COVID-19 patients which manifests as common GI tract symptoms, such as diarrhoea, vomiting, nausea, or abdominal pain (Cheung et al., 2020; Redd et al., 2020; Zhou et al., 2020b).

Since the beginning of the pandemic researchers have carried out sequencing experiments of fecal samples of patients with COVID-19 to uncover a bilateral relationship between COVID-19 and the gut microbiome. According to both alpha and beta diversity indices, SARS-CoV-2 infection was linked to changes in the microbiome community in patients as demonstrated in multiple studies (Kim et al., 2021; Moreira-Rosário et al., 2021; Newsome et al., 2021; Wu et al., 2021; Zhang et al., 2022). Moreover, the Shannon diversity was identified as a risk variable for severe COVID-19 being higher in mild COVID-19 individuals compared to moderate and severe cases. (Moreira-Rosário et al., 2021). Patients hospitalized for COVID-19 have significant changes in stool microbiota composition characterized by an increase in opportunistic pathogens and a decrease in beneficial commensal bacteria compared to controls (Zuo et al., 2020, Moreira-Rosário et al., 2021, Yeoh et al., 2021; Zhang et al., 2022).

There is even more evidence of a change in the taxonomic profile in severely ill patients with COVID-19 compared to healthy or moderately sick patients, but observations might differ in individual studies (Hazan et al., 2022; Sun et al., 2022). Li et al. (Li et al., 2021) discovered that COVID-19 patients had reduced microbial diversity compared to controls, as determined through shotgun metagenomic sequencing and taxonomy indices. Specific bacteria were unique to COVID-19 patients, such as *Streptococcus thermophilus*, *Bacteroides oleiciplenus*, *Fusobacterium ulcerans*, and *Prevotella bivia*. The researchers identified 15 species as microbiological markers for COVID-19 and found relationships between clinical markers and taxonomy. Notably, certain correlations were observed, such as *Coprococcus catus* being positively associated with alanine transaminase levels, red blood cells, and hemoglobin.

Gut microbiome investigations among patients with COVID-19 to date characterized the makeup and diversity of the microbiota through one of two sequencing strategies. Either by targeted amplicon sequencing of a 16S rRNA marker gene (Gu et al., 2020; Tao et al., 2020; Kim et al., 2021; Moreira-Rosário et al., 2021; Newsome et al., 2021; Ward et al., 2021; Wu et al., 2021) or by using deep whole metagenomic (shotgun) strategy (Zuo et al., 2020; Yeoh et al., 2021; Sun et al., 2022). While both strategies are widely used in research, they have limitations in clinical applications of the microbiome as a diagnostic, prognostic, and therapeutic factor in patients with COVID-19. 16S rRNA gene sequencing is a good choice for large sample sizes and cost-efficient analyses, which makes it suitable for use in clinics, however, it has poor taxonomical and functional resolution. On the other side, deep shotgun metagenomics typically costs more but provides greater resolution, allowing a more precise taxonomic and functional classification of sequences (Jovel et al., 2016). The latter, however, is still too costly for all but the most well-funded laboratories and research consortia to implement, creating a potential barrier for diagnostic and prognostic applications that could be adopted by medical and diagnostic facilities. Shallow shotgun sequencing may be a more affordable option than deep shotgun sequencing. It offers nearly the same accuracy at the species and functional level as deep whole metagenome sequencing for known species and genes in five crucial aspects of microbiome analysis — beta diversity, alpha diversity, species composition, functional composition, and clinical biomarker discovery (Hillmann Benjamin et al., 2018).

We conducted an extensive gut microbiome study on 204 hospitalized COVID-19 patients in Poland, employing both shallow and deep shotgun sequencing methods. Our primary objectives were to observe shifts in microbiota composition due to COVID-19 treatment-related hospitalization and associating these changes with clinical factors (e.g., antibiotic use, ICU admission, survival). In comparison to prior studies with smaller cohorts (typically ≤70 subjects, with a maximum of 115), our study featured a significantly larger sample size, allowing for potential confirmation of previous findings.

Additionally, we utilized machine learning techniques to assess the microbiome’s predictive potential for COVID-19 prognosis, comparing its predictive performance with traditional classifiers such as sex, age, body mass index (BMI) and diagnostic findings from laboratory analyses. Notably, we evaluated the utility of shallow shotgun sequencing results as a more cost-effective alternative for clinical diagnostics, benchmarking them against deep shotgun sequencing analysis.

## 2. Materials and methods

### 2.1. Subject recruitment and sample collection

The study group comprised 204 adult patients with confirmed SARS-CoV-2 infection through molecular testing. These patients were hospitalized at the Central Clinical Hospital of the Ministry of Interior and Administration in Warsaw or Teaching Hospital no. 1 Pomeranian Medical University in Szczecin from May 2020 to March 2022. Additional 143 healthy subjects of medical staff working in the hospitals were included as a control group.

Patients were treated according to Evidence Based Medicine and the Polish Ministry of Health treatment guidelines for persons with COVID-19 disease. Exclusion criteria included: lack of consent, a severe clinical condition requiring ICU treatment, and major gastrointestinal and/or abdominal surgery within the last 6 weeks. Demographic, clinical and treatment data, as well as a questionnaire on lifestyle, eating habits, co-morbidities and recent antibiotic therapy, were obtained and managed using REDCap electronic data capture tools (Harris et al., 2009). Stool samples were collected with a swab from faeces gathered on toilet paper into a sterile Eppendorf tube with 2,5 ml ethyl alcohol as preservative and stored at −20°C until DNA extraction. Samples from patients were collected only during hospitalization. A total of 1365 stool samples were gathered, on average 4 (maximum 6) per subject within average 8 days (maximum 70). The study conformed to the Declaration of Helsinki, and all participants signed an informed consent document prior to participation. The study was approved by the institutional review board of the Central Clinical Hospital of the Ministry of Interior and Administration, Warsaw, Poland.

### 2.2. Stool DNA extraction

Nucleic acid extraction was carried out on 942 out of 1365 fecal swabs using the QIAmp PowerFecal Pro DNA kit from Qiagen. Swabs retained for extraction were those that were tightly sealed, ensuring they contained sufficient biological material and preservative inside the tubes. In brief, material from the swabs was transferred into PowerFecal Bead tubes containing buffer C1, followed by homogenization using an Omni Bead Ruptor 12 (with 3 cycles of 30 seconds each, with 30- second breaks in between). Subsequent procedures were conducted following the manufacturer’s instructions. Purified DNA was eluted using 70 µL of the provided elution buffer and quantified using the Quantifluor ONE dsDNA system from Promega.

### 2.3. Shallow shotgun metagenomics sequencing

Sequencing libraries were generated with a reduced volume of KAPA Hyper Plus kit reagent (ROCHE), as described by Sanders et al. in 2019 (Sanders et al., 2019). All steps were carried out in accordance with the manufacturer’s instructions to produce libraries containing metagenomic DNA fragments of approximately 300 bp in size. Initially, metagenomic DNA samples were normalized to a concentration of 10 ng input, followed by a 10-minute enzymatic digestion, indexing with KAPA Unique Dual Indexes (ROCHE), and subjected to 9 cycles of polymerase chain reaction (PCR) library amplification. Subsequently, libraries were purified and size-selected using electrophoretic techniques. The size, quantity, and quality of the selected libraries were assessed using fluorometry with Quantus (Promega) and chip electrophoresis with MultiNA (Shimadzu).

These libraries were further normalized to 2 nM, pooled, denatured with NaOH, and diluted to a final concentration of 8 pM with HT1 buffer (Illumina). These prepared libraries were supplemented with 1% PhiX control v3 (Illumina) and then sequenced on an Illumina MiSeq System using a 2×150-cycles paired-end sequencing strategy, although only the forward reads were used in the subsequent analysis. The Illumina bcl2fastq2 Conversion Software (version 2.20) was employed for demultiplexing sequencing data and converting base call files into FASTQ files using default parameters. On average, 326,385 reads per sample were obtained, with a standard deviation of 93,142.

In the subsequent analyses, 892 samples containing a minimum of 200,000 R1 (forward) reads were included. These analyses encompassed shallow shotgun data profiling, machine learning predictions, and technology comparisons.

### 2.4. Deep shotgun metagenomics sequencing and quality control

Of the samples collected from patients, a subset of 384 samples were selected for deep shotgun sequencing. The same sequencing libraries employed for shallow sequencing were also utilized for deep whole-genome shotgun sequencing of fecal samples, conducted on the Illumina Novaseq6000 platform with a paired-end configuration and a read length of 150 bp. Reads preprocessing was executed using BBTools (BBMap and BBDuk, version 38.96, available at https://sourceforge.net/projects/bbmap/), following the Reads QC Workflow version 1.0.1. This preprocessing involved quality trimming, adapter trimming, and spike-in removal, all carried out using BBDuk. Additionally, human DNA contamination was eliminated using BBMap.

Both the shallow and deep shotgun sequenced data for this study were submitted to the European Nucleotide Archive (ENA) at EMBL-EBI and are accessible under the entry number PRJEB64515.

### 2.5. Shallow shotgun data profiling

Quality control procedures, including the removal of poor-quality reads and adapter trimming (using the adapter sequence ‘AGATCGGAAGAGCACACGTCTGAACTCCAGTCA’), were carried out using fastp (version 0.20.1). The criteria for base qualification were set at a quality value of 15, allowing for a maximum of 40% of unqualified bases. Additionally, a low complexity filter was enabled (Chen et al., 2018). Following quality control, the elimination of human DNA contamination was initially performed by aligning reads to the human reference genome (GRCh38) using minimap2 (version 2.17). Subsequently, reads that did not align were extracted using samtools (version 1.17) (Li, 2018; Danecek et al., 2021). The sequences, now free of contaminants, were aligned to the indexed reference bacterial genome (RefSeq release 82 (O’Leary et al. in 2016)), using Bowtie2. Additional parameters for Bowtie2 were applied: ‘--very-sensitive --no-head --no-unal -k 16 --np 1 -- mp “1,1” --rdg “0,1” --rfg “0,1” --score-min “L,0,-0.05”’. These parameters have been specifically tailored for the purpose of shallow metagenomics, as demonstrated by benchmarking experiments conducted as part of the SHOGUN framework (Hillmann et al., 2020) and subsequently validated by Zhu Qiyun et al. (Zhu Qiyun et al., 2022). Next, we performed operational genomic unit (OGU) profiling using Woltka (https://github.com/qiyunzhu/woltka), obtaining BIOM tables later employed in statistical analyses of shallow shotgun data and machine learning predictions. OGU, a concept similar to the extensively utilized operational taxonomic unit, refers to the smallest unit of microbiome composition that shotgun metagenomic data will permit (Zhu Qiyun et al., 2022). A Github repository for our custom Snakemake (Mölder et al., 2021) pipeline, which implements the methodology described for shallow shotgun sequencing from quality control to classification, is available (https://github.com/bioinf-mcb/polish-microbiome-project/tree/main/shallow-shotgun-analysis-workflow).

### 2.6. Statistical analysis of shallow shotgun data

For shallow shotgun data after rarefying the read count to 100,000 per sample, which left 682 samples, we used QIIME 2 (version 2020.6, (Bolyen et al., 2019)) packages to calculate the alpha diversity (Shannon’s evenness) and beta diversity (weighted UniFrac distance). Weighted UniFrac was selected as our metric because it accounts for both sequence abundance and the relationships among evolutionarily related sequences. To assess the significance of microbial alpha and beta diversities, we employed the Kruskal-Wallis H test and permutational multivariate analysis of variance (PERMANOVA). To examine beta diversity findings, we conducted a principal coordinates analysis (PCoA) on the weighted UniFrac distances within the QIIME 2 framework. To highlight the features (OGUs) with significant effects on the principal component axis, we represented them as arrows in PCoA biplots. To account for changes in the microbiome over time, we conducted pairwise comparisons of beta diversity for samples collected at different time points from the same patient. To further analyze these results in terms of distances from the initial time point and day-to-day changes, we performed linear regressions.

### 2.7. Machine learning predictions

The dataset used to evaluate whether the microbiome can predict COVID-19 outcomes included three types of information: OGUs (obtained from OGU BIOM tables created in Woltka), patient details (like age, sex, and BMI), and clinical test results. We chose this approach to provide the classifier with as much useful information as possible, while minimizing the risk of leaving out important traits. However, including irrelevant or duplicate characteristics could make the classifier overly complex and less able to make accurate predictions for new data. To reduce this risk, we assessed how well the classifier could make accurate predictions by repeatedly testing it with different subsets of the dataset in 51 iterations. To train and evaluate the Random Forest algorithm (Ho, 1995) for disease prediction using microbiota data, we employed a structured approach. We grouped samples by patients to ensure each patient’s data was exclusive to either the training or testing set. In the training set, all available samples from each patient were utilized to enable the algorithm to learn from their microbiota data across different time points, potentially enhancing prediction accuracy.

For the test set, only the initial sample from each patient was used to assess the algorithm’s capability to predict disease based on the patient’s initial microbiome data. The Random Forest algorithm autonomously conducted feature selection by evaluating the importance of each feature in predicting the target variable (ICU admission/death). Feature importance scores were determined using the mean decrease impurity measure, which quantifies a feature’s contribution to reducing impurity, as measured by the Gini index, in the decision trees of the Random Forest.

We employed AUC-ROC as an evaluation metric to gauge the random forest classifier’s performance. It illustrates the classifier’s ability to discriminate between positive and negative samples by plotting sensitivity against 1-specificity at different thresholds. The AUC-ROC score ranges from 0.5 (random guessing) to 1 (perfect classification), with higher values denoting better performance.

### 2.8. Comparison of shallow and deep shotgun data

The dataset for this section comprised sequenced samples from COVID-19 patients that had undergone quality control procedures, as previously described, and represented the intersection of data obtained through both shallow and deep whole metagenome approaches (193 samples). Control samples were deliberately omitted from the dataset, as the objective of the analysis centered on the assessment of employing shallow sequencing in lieu of deep sequencing for discerning COVID-19-associated microbiome modifications. To maintain consistency, all samples in the dataset were profiled using Metaphlan4 (Blanco-Míguez et al. 2023) with default settings. The comparison of shallow and deep sequencing was performed using QIIME 2 (Bolyen et al., 2019) or custom Python scripts. Alpha and beta diversities were compared using QIIME 2 diversity modules, and the metrics used were Shannon entropy, observed features (alpha diversity) and Bray-Curtis dissimilarity (beta diversity).

## 3. Results

### 3.1. Demographic and clinical characteristics of the study’s subjects

Table 1 lists the demographic and clinical characteristics of patients (n = 204) and controls (n = 143). All patients were Polish residents. Out of 204 patients, men made up 125 (61.3%), and women 79 (38.7%). The mean ± standard error of the mean age in years was 61.2 ± 1.3 (range, 17–96) for patients. A large majority of patients overall (202 or 99.0%) were White. One patient was Latino and one patient was mixed-race. Antibiotics were administered to 57.4% of the SARS-CoV-2-infected patients. In terms of the outcome, 170 patients (83.3%) were released from the hospital, and 34 (16.7%) died because of COVID-19 while they were in the hospital. Additionally, 50 patients (24.5%) were admitted to the ICU, and 154 (75.5%) patients were continuously hospitalized in the dedicated COVID-19 unit.

**Table 1.** Summary of the COVID-19 patients from the study cohort.

|  | **COVID-19 patients** |  |
| --- | --- | --- |
|  | **Before rarefaction** | **After rarefaction to 100, 000 features per sample** |
| **Number of participants** | 204 | 176 |
| **Age, mean years** | 61.2 | 61.2 |
| **Sex** |  |  |
| Male (%) | 125 (61.3) | 103 (58.5) |
| Female (%) | 79 (38.7) | 68 (41.5) |
| **Ethnicity** |  |  |
| White (%) | 202 (99.0) | 174 (98.8) |
| Latino (%) | 1 (0.5) | 1 (0.6) |
| Mixed (%) | 1 (0.5) | 1 (0.6) |
| **Hospitalization outcome** |  |  |
| Death (%) | 34 (16.7) | 28 (15.9) |
| Survival (%) | 170 (83.3) | 148 (84.1) |
| **Antibiotics treatment during hospitalization** |  |  |
| Yes (%) | 117 (57.4) | 98 (55.7) |
| No (%) | 87 (42.6) | 78 (44.3) |
| **ICU referral** |  |  |
| Yes (%) | 50 (24.5) | 41 (23.3) |
| No (%) | 154 (75.5) | 134 (76.7) |

### 3.2. The gut microbiome of COVID patients differs from that of non-COVID controls

We examined changes in the fecal microbial composition of actively infected SARS-CoV-2 patients over time by comparing weighted UniFrac distances between a patient’s (case) initial and subsequent sample points, in contrast to the control group. Interestingly, the distance between control samples remained relatively stable over time, while the distance between patient samples increased as time progressed (Fig. 1A). Furthermore, in our comparison of samples on a day-to-day basis, we observed that the distances were more substantial for the patient group and exhibited a slower rate of decrease compared to the control group (Fig. 1B).

**Figure 1.**
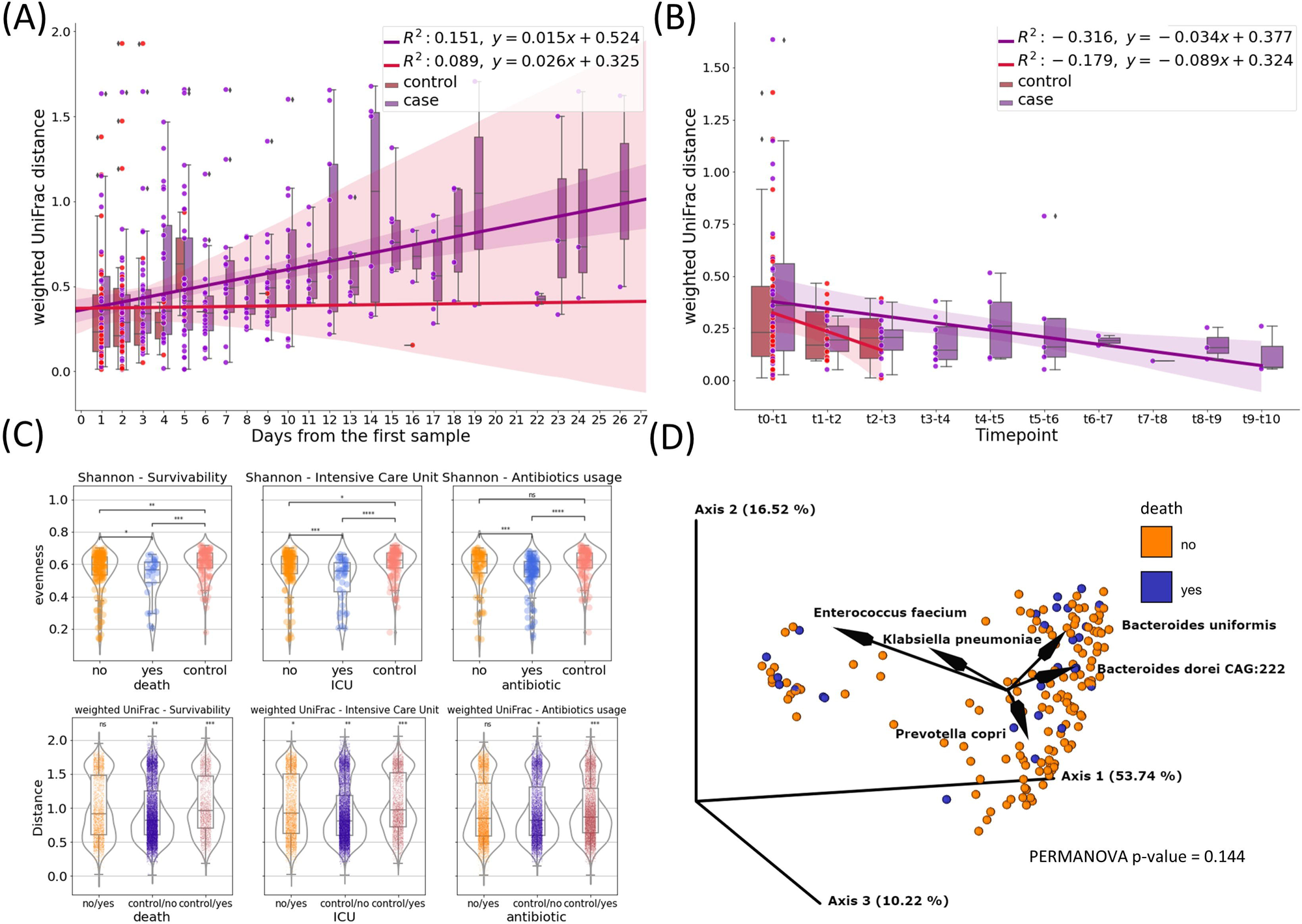
The gut microbiome of COVID patients. (A) Subject’s weighted UniFrac distances to subject’s first sample after rarefaction, showing how the composition of fecal microbes in SARS-CoV-2 patients change over time compared to the control group. Linear regression models the relationship between distance and time. (B) Subject’s day-to-day change of weighted UniFrac distance. Linear regression shows the correlation between the distance and time point. (C) Microbiome diversity measures - Shannon’s entropy and weighted UniFrac for survivability, Intensive Care Unit referral and antibiotics usage measured for subject’s first sample after rarefaction (death, icu) or first sample collected after antibiotic introduction (antibiotics) (ns - not significant; * - 0.01 < p ≤ 0.05; ** - 0.001 < p ≤ 0.01; *** - 0.0001 < p ≤ 0.001; **** - p ≤ 0.0001) (D) PCoA biplot of weighted UniFrac of subject’s oldest sample after rarefaction coloured by survivability with taxa contributing to the PCoA axes.

We compared the microbiome diversity of patients based on their hospitalization outcomes (survival or death), ICU referral status (yes or no), and antibiotic treatment (treated or untreated) using their earliest or post-antibiotic introduction samples, while also including control samples as a separate category. According to Shannon’s evenness analysis, patients who passed away due to COVID-19 differed significantly from those who recovered (p ≤ 0.05). The difference was more pronounced when comparing surviving patients to controls (p ≤ 0.01), and most significant when contrasting deceased patients with healthy controls (p ≤ 0.001). While no statistically significant difference in weighted UniFrac was observed between surviving and non-surviving patients in pairwise comparisons of beta diversity distances based on hospital outcomes, a level of significance was detected when comparing patients to controls (p-value for surviving patients vs. controls, p ≤ 0.01; dead patients vs. controls, p ≤ 0.001).

In terms of ICU admission, the most significant diversity variations were observed between patients referred to the ICU and those solely in the COVID unit, as well as between ICU-referred patients and controls (p ≤ 0.0001 to 0.001). Regardless of hospitalization type, ICU referral consistently led to statistically significant differences in weighted UniFrac distances, particularly when compared to controls.

In contrast, patients not treated with antibiotics showed similar diversity levels as the control group, while those receiving antibiotics exhibited higher Shannon diversity (p ≤ 0.0001 to 0.001) compared to both untreated patients and controls. The difference was most pronounced in patients who received antibiotics (p ≤ 0.001). However, there were no significant variations in beta diversity between treated and untreated patients. (Fig. 1C).

We employed weighted UniFrac-based principal coordinate analysis and looked for any metadata variables that could explain the behavior of the data points on the PCoA plot (Fig. 1D). 80.48% of the total variation in the SARS-Cov-2 patients was described by the first three PCoA components (ie. PC1-PC3). We were unable to identify a single demographic or clinical variable that would explain the distribution, but by creating a PCoA biplot, we were able to determine which taxa contribute the most to the PCoA axes (Fig 1D). The presence of *Enterococcus faecium* in the patient samples accounts for the major variation. The remaining four OGUs - *Bacteroides uniformis*, *Klebsiella pneuomoniae*, *Bacteroides doreii CAG:222*, and *Prevotella copri*, are also largely responsible for the divergence.

### 3.3. Machine learning predictions

Our objective was to ascertain the most critical information for accurately predicting patient outcomes. To accomplish this, we devised six distinct classifiers, each designed to analyze different sets of input data: baseline, clinical, metadata, microbiome, microbiome combined with clinical, and microbiome combined with metadata.

Regarding the prognosis of ICU admission, all classifiers outperformed the baseline significantly based on the ROC-AUC score (Fig. 2A). To further assess and compare these classifiers, we conducted ANOVA analysis, revealing that their performance was strongly influenced by the availability of features. Microbiome-based classifiers demonstrated the highest performance, and the inclusion of additional data, whether clinical or metadata, did not provide a substantial advantage. In contrast, classifiers that did not utilize microbiome data performed notably worse, with metadata-based classifiers showing only marginal improvement over the baseline.

**Figure 2.**
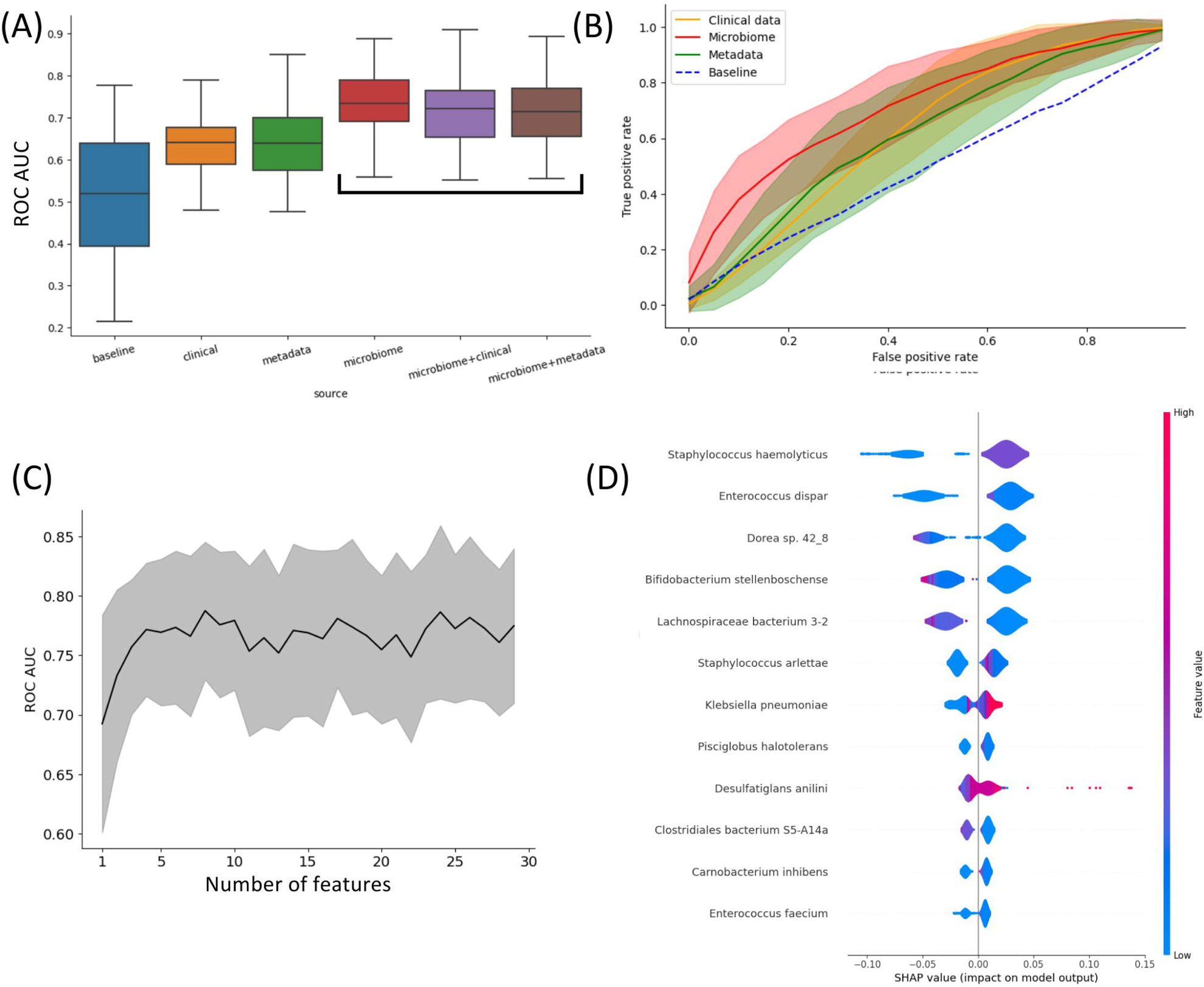
Insights into what influences the predictive power of patients’ outcomes (ICU vs non-ICU) classifier. (A) Impact of different types of data on the predictive power of the classifiers. This plot shows that access to microbiome data immensely increases the performance of the classifiers. (B) ROC curve of classifiers grouped by access to data. (C) Increasing the number of metagenomic features doesn’t improve ROC-AUC beyond the 7 most important. (D) Shapley values of the most important features for classification.

In Fig. 2B, the ROC curves of the four main classifier types (clinical, metadata, microbiome, and baseline) are compared, highlighting that the microbiome classifier’s enhanced AUC is primarily attributed to its ability to achieve a significantly higher True Positive Rate for small False Positive Rates compared to other classifiers.

Remarkably, only four features (taxa) are necessary to achieve optimal performance for the microbiome-based classifier (Fig. 2C). The assessment of feature importance revealed that high concentrations of *Orrella dioscoreae* and *Klebsiella pneumoniae* correlated with worse outcomes, while the presence of *Lachnospiraceae bacterium 3-2* was associated with improved patient prognosis (Fig. 2D).

A comparable analysis of the life/death outcome is available in the supplementary material (Fig. S1).

### 3.4. Shallow vs deep shotgun comparison

To validate the suitability of using shallow sequencing instead of deep shotgun sequencing in COVID-19 patients, we conducted a comparative analysis of matched samples from our study. Shallow and deep sequencing samples exhibited no significant differences in fundamental quality parameters such as read length or GC content. The relatively higher rate of quality control failures in deep sequencing reads could be attributed, in part, to a greater duplication rate compared to shallow sequencing (Fig. S2). While alpha diversity and some observed features were higher in deep sequencing, there was no distinct separation between the two sequencing types when performing beta diversity clustering (Fig. 3A, Fig. S3).

**Figure 3.**
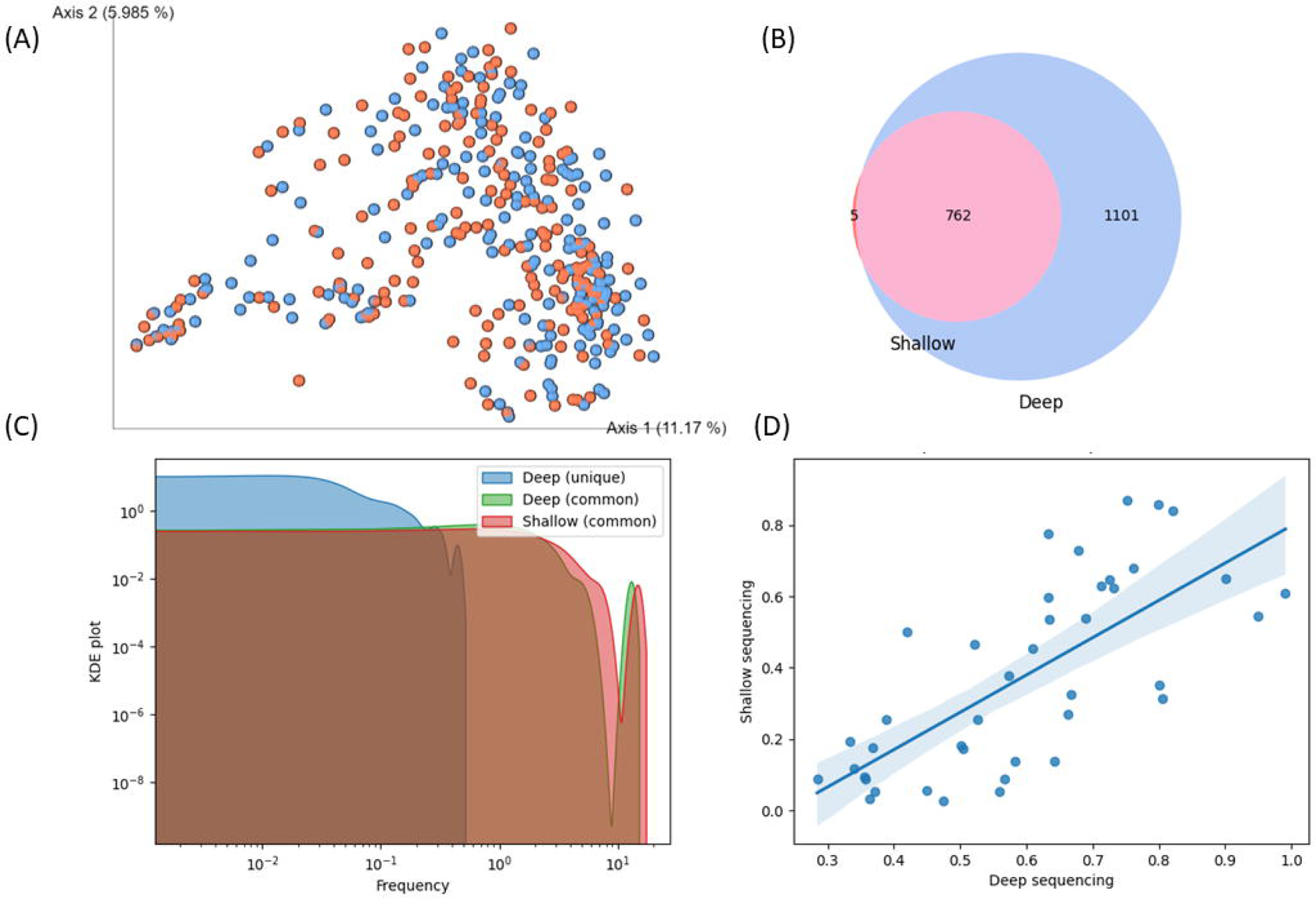
Comparison of shallow and deep shotgun sequencing methods. (A) Clustering of samples based on Bray-Curtis beta diversity. Blue: deep, red: shallow sequencing. (B) Overlap of species identified in shallow and deep sequencing. (C) KDE plot of species abundance identified uniquely or commonly in shallow and deep sequencing. (D) Correlation of species abundance in shallow and deep sequencing, restricted to abundance below 1% in an exemplary sample. Spearman = 0.75, p-value = 0.0, Mean squared error = 9.04, R-squared = 0.71.

A high degree of overlap of species identified in shallow and deep sequencing was observed (Fig. 3B). While substantially more species were found in deep sequencing, all but five species identified in shallow sequencing were discovered in deep sequencing. The five species unique to shallow sequencing were *CAG-269_sp900554175*, *Faeciplasma gallinarum*, *Klebsiella pneumoniae*, *Mediterraneibacter glycyrrhizinilyticus A*, *Parafannyhessea umbonate* and *Scatacola A* faecigallinarum. They rarely appear in bioinformatics analyses, and Klebsiella pneumoniae is known to be frequently misclassified (Arnold et al., 2011). Species identified in both shallow and deep sequencing had an abundance of at least 0.2% in deep sequencing. Any species below this threshold were not detected in shallow sequencing (Fig. 3C). Considering that most studies determine 1% as a cut-off (Cena et al., 2021), our result indicated a similarity between the range of species detected by both methods. In addition to that, we showed that while the abundance of species was not perfectly matched between shallow and deep sequencing, the hierarchy of species abundance, even below 1%, was well maintained (Fig. 3D).

## 4. Discussion

In the wake of the COVID-19 pandemic, the scientific community has devoted significant effort towards investigating the pathogenesis of SARS-CoV-2 infection and identifying the risk factors that contribute to disease outcomes. As part of these efforts, our study explored the potential role of gut microbiota as a risk factor for ICU referral or mortality in individuals with COVID-19. Using both shallow and deep sequencing techniques, we studied the gut microbiomes of 204 COVID-19 patients at two reference hospitals in Poland. We sought to learn how hospitalization affected the makeup of the microbiota and how these changes related to patient outcomes. The study employed machine learning to see if microbiome data might predict COVID-19 prognosis more accurately than conventional predictors like age, sex, and BMI. Using both shallow and deep sequencing techniques allowed us to contrast their precision, specifically to find out if shallow sequencing can serve as a potential cost-effective substitute with excellent taxonomic accuracy for COVID-19 patient clinical outcomes prediction.

The fecal microbial beta diversity of the SARS-CoV-2 patients who are actively infected increases over time as compared to that of the hospital staff, whose distance almost remains constant over time (Fig. 1). Additionally, day-to-day comparisons revealed that the distances are greater and are shrinking more slowly for the patients than for the control group. This suggests that the microbiome of COVID-19 hospitalized patients is less stable and subject to greater qualitative and quantitative perturbations over time compared to healthy controls.

We were able to distinguish patients stratified by survivability from healthy subjects when both alpha (Shannon’s evenness) and beta (unweighted UniFrac) heterogeneity were compared, as the differences between these groups were significant in both cases (Fig. 1). The highest significance was observed for deceased patients matched against controls. Similarly, the metrics of both diversities, alpha and beta, are most important for the patients admitted to ICU paired with controls. It should be noted that although the difference was smaller, we also observe a significant difference between patients who only stayed in the COVID-19 ward and those who were referred to ICU.

Most of the variation in the unweighted UniFrac PCoA plot can be attributed to the presence of *Enterococcus faecium* in patient samples. The plot’s divergence is also largely attributable to the other four OGUs, *Bacteroides uniformis*, *Klebsiella pneuomoniae*, *Bacteroides dorei CAG:222*, and *Prevotella copri*.

Using patient metadata, microbiome and clinical data, we carried out an in-depth machine-learning analysis. Our findings shed light on the varying impacts of different combinations of clinical, microbiome, and patient metadata on the accuracy of outcome prediction for patients and suggest that the AUC-ROC of the classifiers is primarily influenced by their access to microbiological data, indicating that microbiological data is a more reliable predictor of patient outcomes compared to clinical or metadata. Our analysis of feature importance additionally proves that only a few of the taxa are important in the prediction of patients’ outcomes.

However, our results do not allow us to conclude unequivocally that the observed dysbiosis is a causal factor for the severe course of the disease or a consequence of it. Gastrointestinal dysbiosis in COVID-19 can occur due to antibiotic therapy, secondary bacterial infections, and enteral nutrition (Langford et al., 2020; Zaher, 2020). Altered microbiota can cause inflammation in the gastrointestinal tract, malnutrition (Zaher, 2020), and viral and bacterial infections (Zuo et al., 2020). COVID-19 patients can also have an altered gut microbiota before the disease and/or hospital admission (Alberca et al., 2021). In these patients, COVID-19 may exacerbate dysbiosis leading to different health complications like metabolic disturbances (Alberca et al., 2021).

We have proven that shallow shotgun sequencing is a valid alternative to deep sequencing for predicting COVID-19. Although deep sequencing detected more species and had higher alpha diversity, there was no significant difference in beta diversity clustering between the two methods. The range of species detected by both methods was similar, and the abundance of species was maintained in a proper hierarchy. Our findings suggest that shallow sequencing may be a viable substitute for deep sequencing in clinical settings. Shallow shotgun sequencing has been demonstrated to yield quicker findings in a clinical context, and it also offers better economic viability when used with popular and widely accessible Illumina platforms like MiSeq. Shallow shotgun sequencing, which is substantially less expensive than deep shotgun sequencing, provided lower technical variation and higher taxonomic resolution than 16S sequencing, according to Reau et al. (La Reau et al., 2023). As bioinformatics techniques are developed and standardized and computational performance increases, the use of in situ microbiome characterization in the therapeutic context is becoming more and more accepted.

## Abbreviations

AUC-ROC: area under the receiver operating characteristic curve
BMI: body mass index
COVID-19: Coronavirus Disease 2019
GI: gastrointestinal
ICU: intensive care unit
MAG: metagenome-assembled genome
OGU: operational genomic unit
PCoA: principal coordinates analysis

---

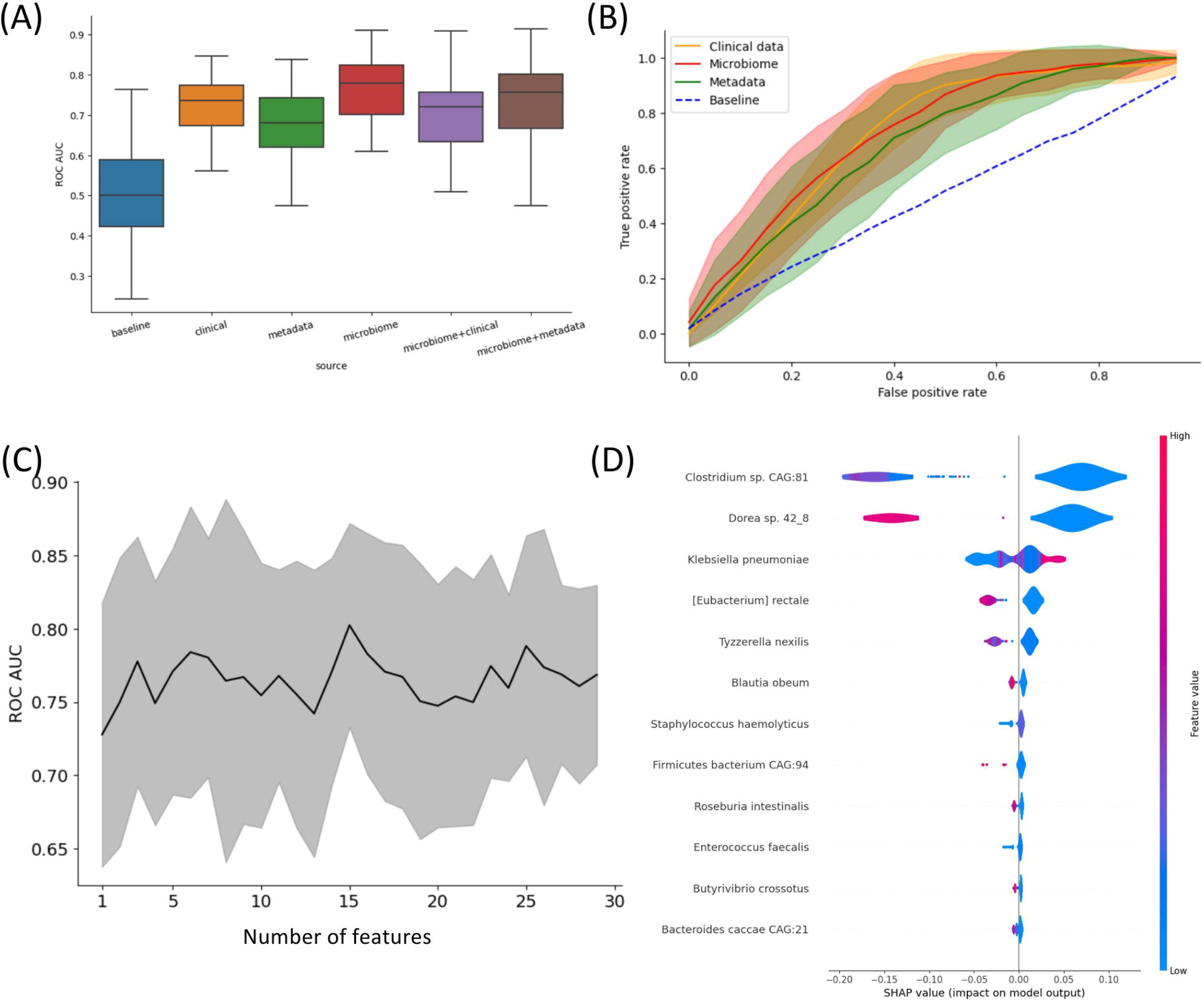

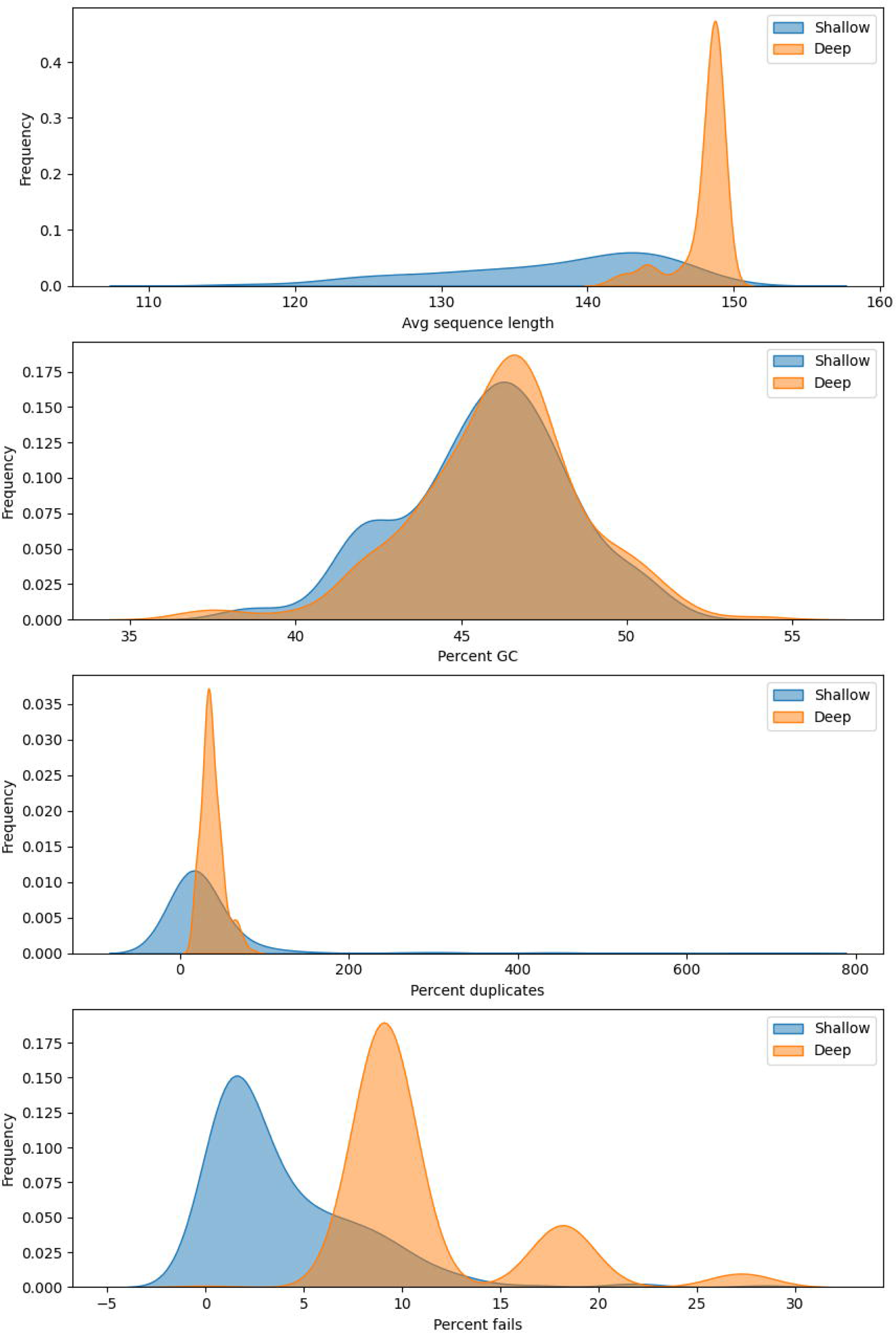

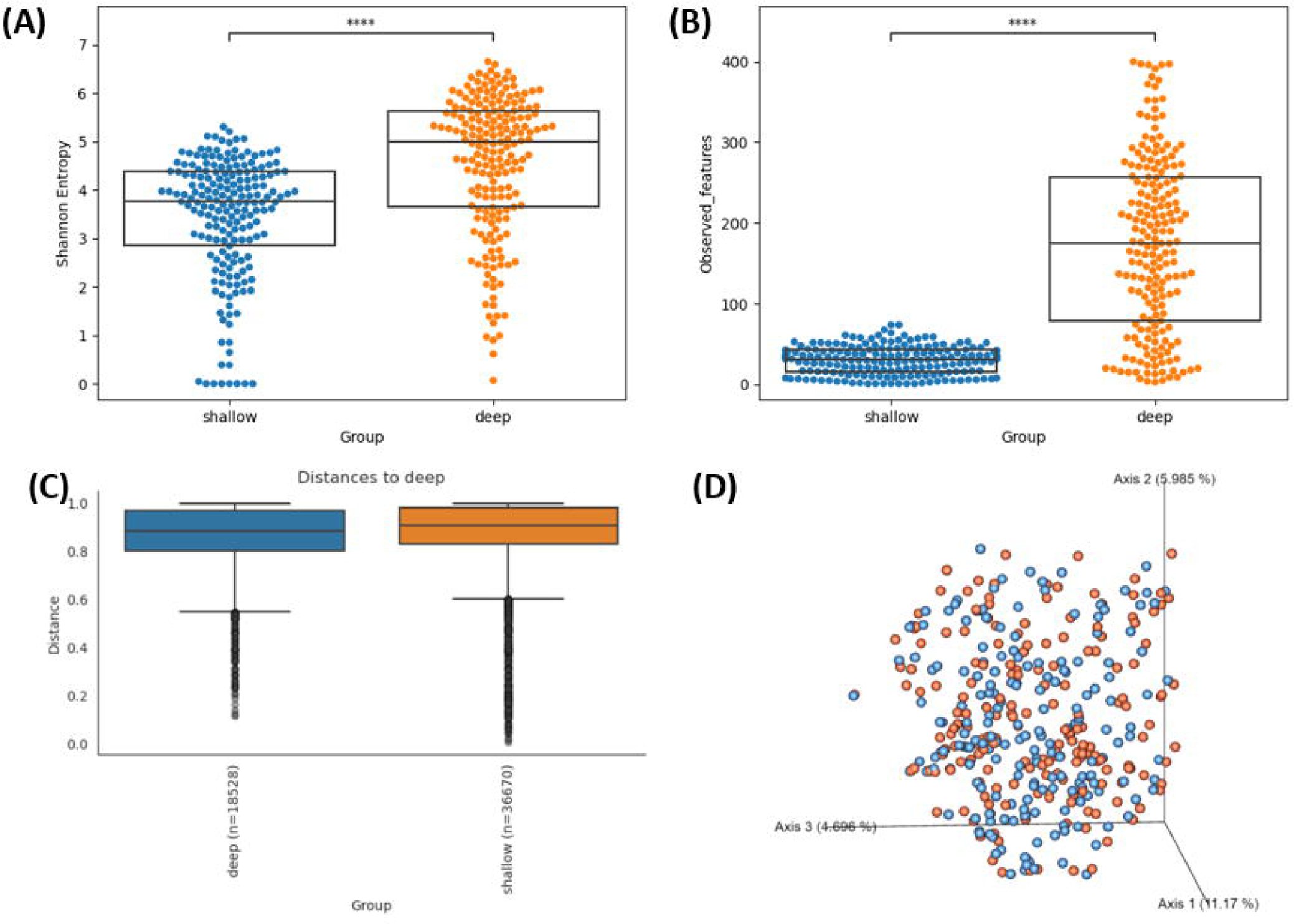

