## Supplementary material for "Gut Microbiome Dynamics and Predictive Value in Hospitalized COVID-19 Patients: A Comparative Analysis of Shallow and Deep Shotgun Sequencing"

### 1. Supplementary Figures

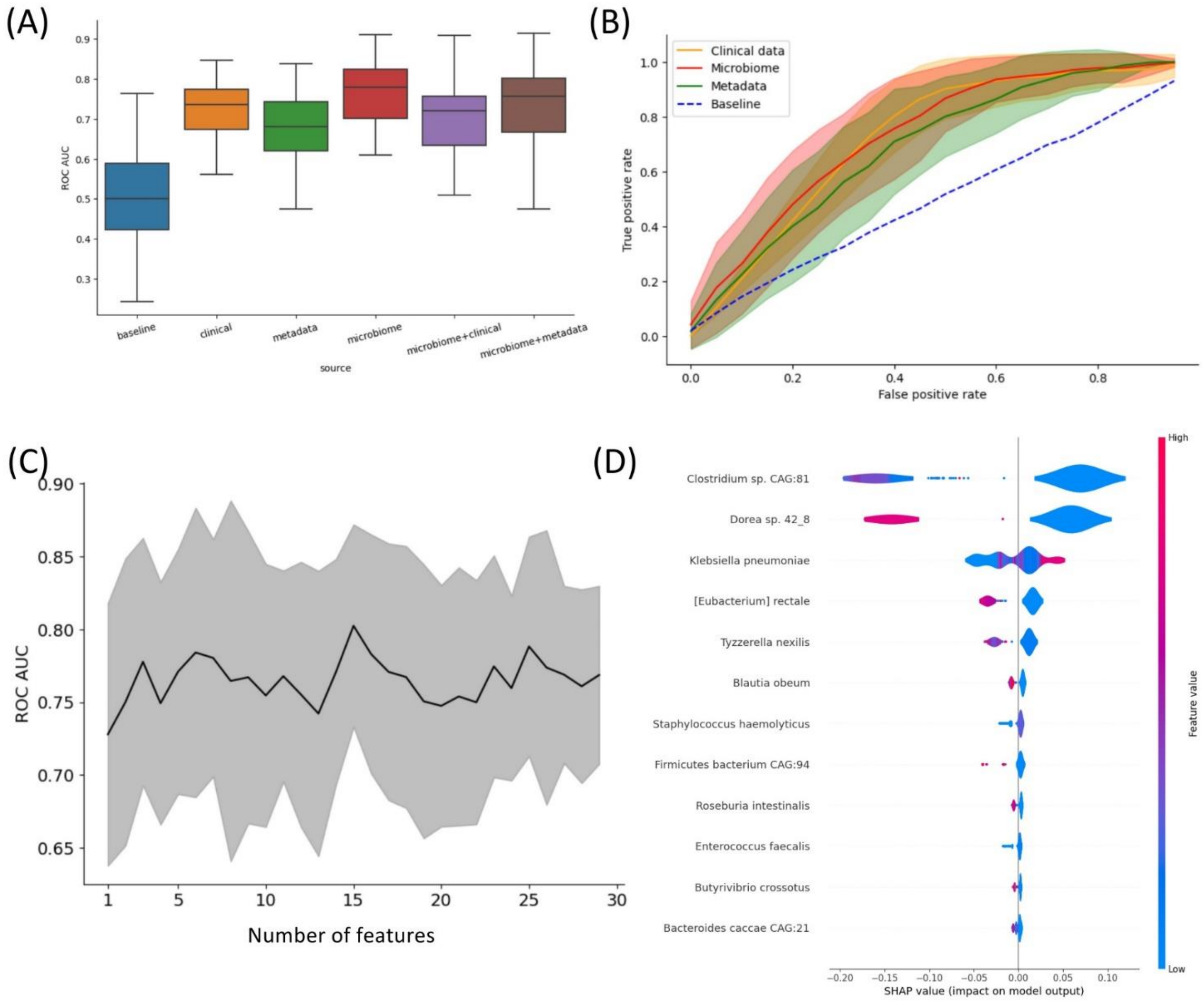


**Supplementary Figure 1. Insights into what influences the predictive power of patients' outcomes (life vs death) classifier.** (A) Impact of different types of data on predictive power of the classifiers. This plot shows that access to microbiome data increases the performance of the classifiers. The three underlined classifiers form a cluster with no inter-cluster difference. (B) ROC curve of classifiers grouped by access to data. (C) Increasing number of metagenomical features doesn't improve ROC-AUC beyond the first most important taxon. (D) Shapley values of the most important features for classification. Klebsiella pneumoniae is strongly connected to increased chance of patient's death, as well as staphylococcus haemolyticus.

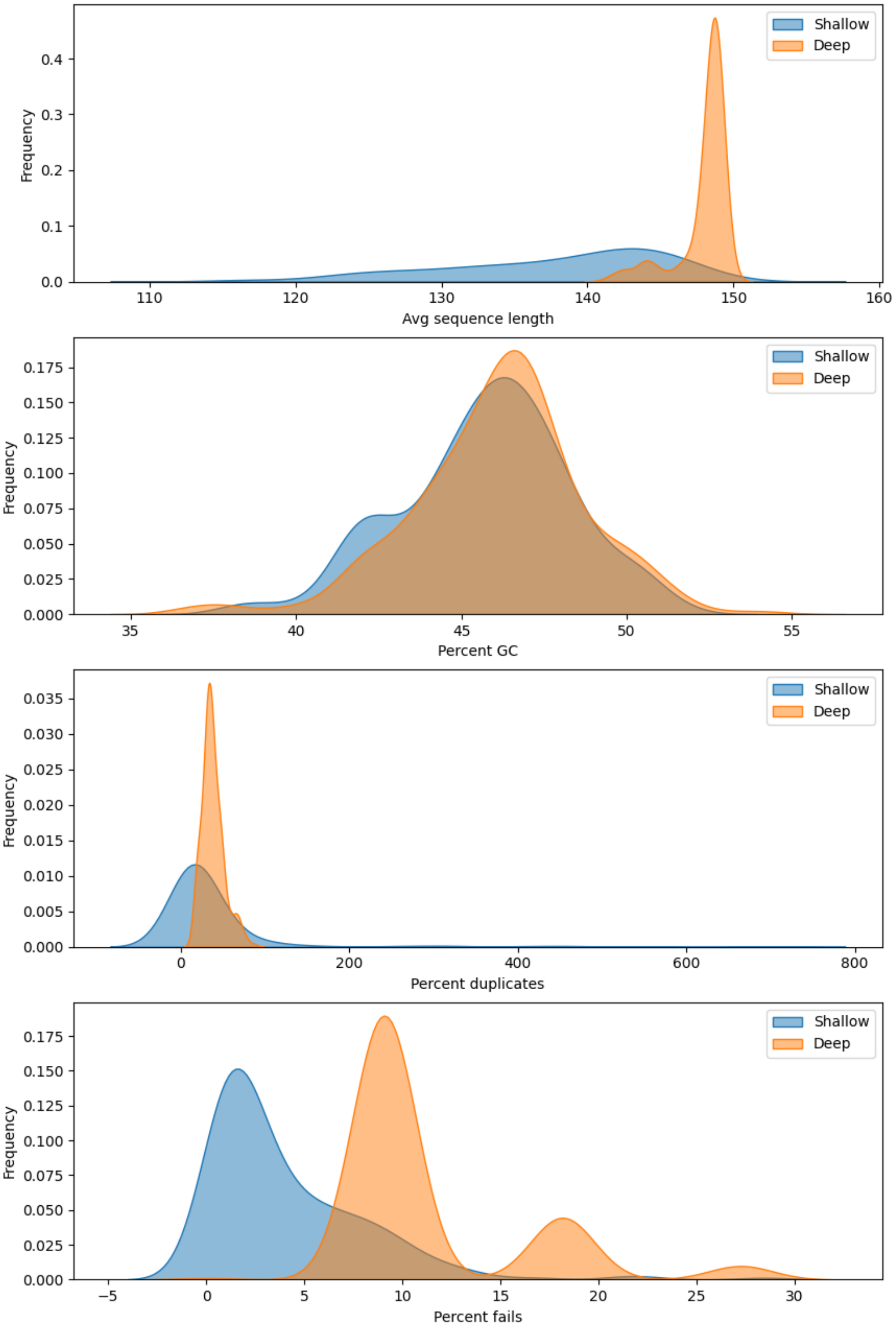


**Supplementary Figure 2. Quality comparison of matched shallow and deep sequencing samples.**

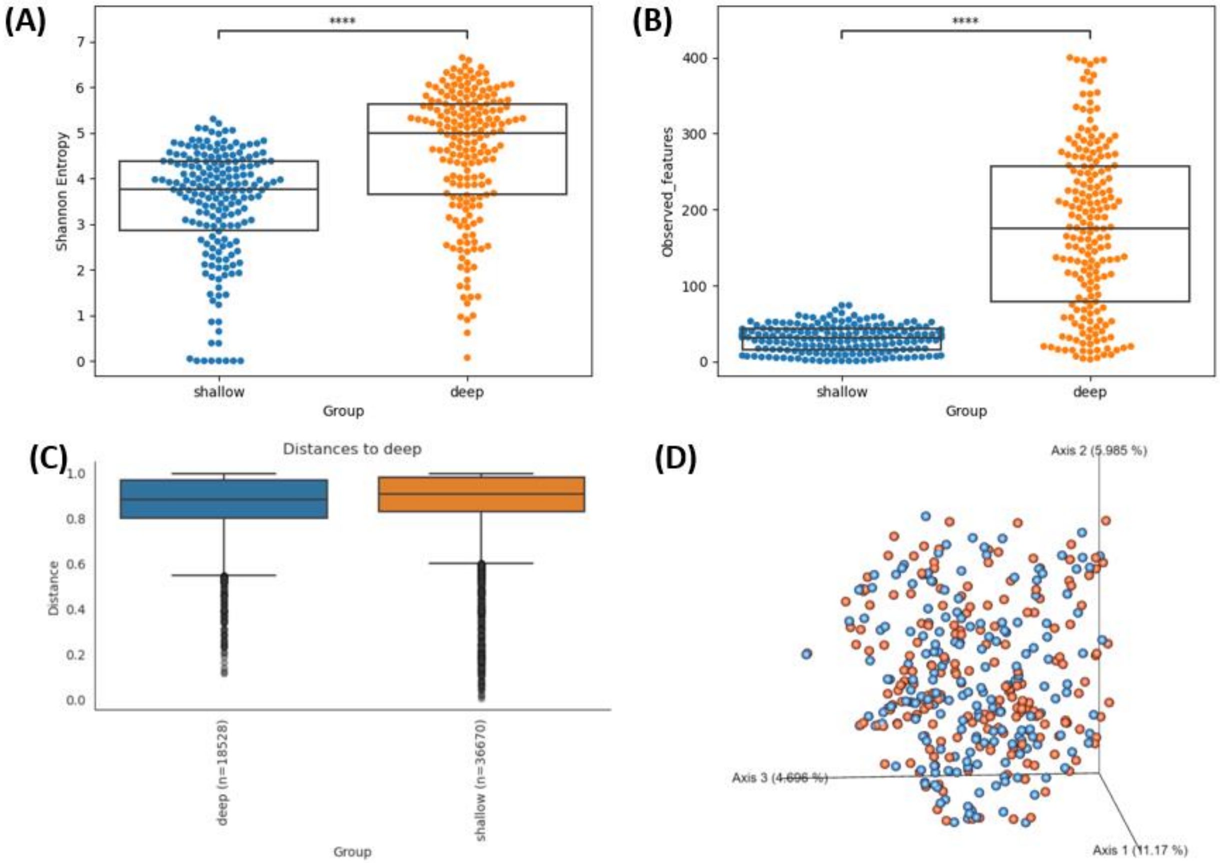


**Supplementary Figure 3. Comparison of shallow and deep sequencing samples in terms of a) alpha diversity (Shannon entropy) and b) number of observed features. c) Bray-Curtis beta diversity. d) Emperor plot of Bray-Curtis beta diversity.** Red: shallow, blue: deep sequencing.
